# Population-scale transcriptomics reveals host genetic control of phyllosphere fungal communities

**DOI:** 10.64898/2026.05.21.727028

**Authors:** Charles Colvin, Surinder Chopra

## Abstract

The composition of plant-associated microbial communities plays a central role in host health and agricultural productivity, yet the genetic basis of plant–fungal interactions in the phyllosphere remains poorly understood. Progress has been limited in part by technical challenges in profiling low-abundance fungal communities and linking their variation to host genetics. Here we show that standard polyA-enriched RNA sequencing of plant leaf tissue, without targeted microbial enrichment, captures sufficient fungal transcripts to enable quantitative profiling of metabolically active mycobiomes at scale.

Although fungal transcripts represented only 0.03–0.4% of classified reads in standard polyA-enriched leaf RNA-seq datasets, this still yielded >79 million fungal reads across 2,194 field-collected samples (median = 5,260–59,746 reads per sample), enabling robust quantification of several hundred fungal taxa. Leveraging these data, we perform genome-wide association and transcriptome-wide association analyses of fungal relative abundance and identify widespread host genetic control of phyllosphere fungal communities.

We detect extensive and biologically structured associations between host genetic variation, gene expression, and fungal abundance across all three crops, including numerous loci with multi-taxon effects and strong colocalization between GWAS and expression quantitative trait loci. Network and module-level analyses further reveal coordinated host transcriptional programs linked to fungal community variation, while integration of orthologous gene sets indicates partial conservation but substantial species specificity in underlying genetic architecture.

Together, these results demonstrate that standard RNA-seq datasets contain sufficient ecological signal to enable high-resolution microbiome genetic mapping and reveal that host genetic control of phyllosphere fungal communities is more pervasive than previously appreciated. This framework provides a scalable approach for reusing existing transcriptomic datasets to study plant–microbe interactions across diverse species and environments.

## Introduction

Plant-associated microbial communities consist of beneficial endophytes, growth-enhancing bacteria, nitrogen-fixing microbes, mycorrhizal fungi, and other symbiotic organisms. These communities also comprise plant pathogens and a range of commensal or transient taxa that may either lack obvious functional roles or serve as environmental contaminants from soil, air, or surface interactions. Understanding the composition and dynamics of these communities serves multiple research objectives: discovery of microorganisms with agricultural potential such as biofertilizers and plant growth-promoting agents, identification of antagonistic interactions that could be leveraged for biological disease control [1], and elucidation of plant genetic factors governing microbial colonization that may inform breeding strategies for pathogen resistance. 5-7% of annual losses in U.S. maize production, equivalent to 1.1-1.8 billion bushels are lost to disease each year, with the majority of losses resulting from fungal pathogens [2]. Moreover, the ongoing evolution of pathogen virulence and breakdown of known resistance loci underscores the need for deeper mechanistic understanding of plant-microbe interactions.

Recent technological advances in high-throughput sequencing have enabled comprehensive characterization of bacterial and fungal communities across distinct plant compartments: the rhizosphere (root-soil interface), endosphere (internal tissues), and phyllosphere (on/within aerial tissues) [3,4]. These profiling efforts have revealed that community structure varies both with environmental conditions, even among genetically uniform plants, and with host genotype when comparing genetically diverse individuals in shared environments [3–6]. The proportion of microbial variation attributable to plant genetics varies considerably across studies, with ongoing debate regarding whether foliar or root-associated communities show stronger host genetic influence [3–5,7–10].

Genome-wide association mapping has emerged as a powerful approach to connect host genetic variation with microbial abundance patterns. Prior investigations have successfully identified loci associated with variation for both bacterial and fungal communities in Arabidopsis leaves across field environments [7,11], maize foliage [9], and root-associated bacterial communities in sorghum [5], maize [10], and Brassica napus [12]. The latter study additionally employed transcriptome-wide association analysis to pinpoint specific genes whose expression correlates with the relative abundance of multiple root bacteria. However, results from leaf microbiome studies have been mixed: an eight-environment Arabidopsis study detected only two associations surpassing stringent multiple-testing thresholds [11], while work on maize leaf bacteria identified few significant loci and attributed most variation to environmental factors and stochastic colonization events [9].

Amplicon-based sequencing remains the dominant methodology for bacterial and fungal community profiling. This approach employs primers targeting conserved genomic regions that flank variable sequences, generating short amplicons that can be sequenced efficiently across many samples. While cost-effective and analytically streamlined, amplicon methods present several limitations: primer-dependent taxonomic biases leading to uneven representation, insufficient resolution for discriminating closely related organisms, and extensive zero-inflation arising from shallow sequencing depth combined with numerous PCR cycles [13,14]. Furthermore, standard DNA-based protocols cannot distinguish metabolically active cells from dormant spores or residual DNA from dead organisms, potentially distorting community composition estimates [15]. RNA-based approaches targeting the same amplicons have demonstrated improved associations with host genetics compared to DNA-based methods [16,17], though they do not address PCR-related artifacts.

Transcriptomic profiling of total community RNA offers a compelling alternative that simultaneously addresses the viability problem, capturing only metabolically active organisms and minimizing PCR amplification artifacts inherent to amplicon workflows. However, the substantially higher sequencing requirements and library preparation costs have historically limited the scale of such studies, precluding the large sample sizes necessary for robust genetic association analyses.

Here we demonstrate that polyadenylated RNA-sequencing datasets generated from field-collected leaf tissue yield sufficient fungal-derived reads for community profiling without specialized mycological protocols. Leveraging existing population-scale RNA-seq datasets from maize, sorghum, and soybean spanning three distinct environments and encompassing more than 2190 total samples [18–20], we address three central questions: (1) How does phyllosphere fungal community structure vary among crop species and across environments? (2) To what extent do host genetic factors shape fungal community composition, and which plant genes, pathways, or regulatory modules are most strongly associated with fungal abundance? (3) Can we integrate genome-wide genetic mapping with transcriptomic association signals to identify specific host loci controlling fungal colonization?

## Methods

### RNA-seq source, processing and taxonomic assignment

We obtained raw RNA-seq data from three separate, previously published studies of population-scale transcriptomics in field environments consisting of sorghum (*Sorghum bicolor*) [18], maize (*Zea mays*) [19], and soybean (*Glycine max*) [20]. Raw sequencing data for these studies are publicly available through the European Nucleotide Archive (ENA) under accession numbers PRJEB83049 (sorghum) and PRJEB67964 (maize), and through the Genome Sequence Archive (GSA, Beijing Institute of Genomics Data Center) under accession CRA009979 (soybean). The details for germplasm selection, plant growth, sample collection and library preparation have been described previously in the original studies.

Briefly, the sorghum dataset consists of 822 libraries drawn from a superset of the Sorghum Association Panel [21] and the Sorghum Diversity Panel [22] grown together in Lincoln, NE, USA in 2021. The maize dataset contains 750 libraries from the Wisconsin Diversity (WiDiv) panel [23] grown in Lincoln, NE, USA in 2020. The soybean dataset consists of 622 libraries collected from a diversity panel grown in Sanya, China.

For maize and sorghum, samples were obtained from leaf punches collected from the pre-antepenultimate leaf (the fourth leaf from the flag leaf, if present, or the topmost visible and fully emerged leaf if no flag leaf was present). In contrast, soybean samples were collected from all tissues above the cotyledon node at the V2 developmental stage. All samples within each dataset were collected within a narrow time window on a single day to minimize temporal variation.

For maize and sorghum, genetic marker data were obtained from publicly available repositories (Dryad: https://doi.org/10.5061/dryad.bnzs7h4f1 and Figshare: https://doi.org/10.6084/m9.figshare.27936195, respectively). The maize marker set was derived from whole-genome resequencing [24] and includes 696 individuals overlapping with the RNA-seq dataset. The sorghum variant dataset, derived from RNA-seq-based variant calling as described in the original study [18], includes 736 individuals with matching expression data.

Publicly available variant data were not available for the soybean population; therefore, soybean samples were excluded from analyses requiring genotype information, including genome-wide association studies (GWAS), expression quantitative trait locus (eQTL) mapping, and narrow-sense heritability estimation.

Raw RNA-seq reads were trimmed to remove low quality and adapter sequences using Trimmomatic (v0.33) [25] with the parameters ILLUMINACLIP:TruSeq3-PE.fa:2:30:10 LEADING:3 TRAILING:3 SLIDINGWINDOW:4:15 MINLEN:35. Taxonomic classification of trimmed reads was performed using Kraken2 (v2.1) [26] with the prebuilt *core_nt* database, a curated subset of the NCBI nucleotide (nt) database [27], including bacterial, archaeal, viral, fungal, and other eukaryotic reference sequences (version 10/15/2025; downloaded from https://benlangmead.github.io/aws-indexes/k2). Taxonomic abundance was estimated at five taxonomic ranks: class (C), order (O), family (F), genus (G), and species (S), from the Kraken reports and *core_nt* database using Bracken (version 3.1) [28]. For estimating relative fungal abundance, only reads assigned to taxa classified as fungal (kingdom Fungi, NCBI:txid4751) were considered, and all proportions were calculated relative to total Bracken-classified reads at the given taxonomic level.

Rarefaction curves were generated at all five taxonomic ranks using the vegan R package (version 2.7-1) [29]. After retaining only those fungal taxa whose presence was supported by at least 25 RNA-seq reads in at least one third of all samples from a given dataset, alpha diversity metrics were calculated using the phyloseq package (version 1.46.0) [30,31]. Fungal co-occurrence networks were estimated from filtered count data using the Spiec-Easi R package (version 1.1.3) [32] with the neighborhood selection method (method = ‘mb’), lambda.min.ratio = 1×10⁻², nlambda = 30, and stability selection via the pulsar algorithm (rep.num = 100).

### Filtering and normalization of fungal abundance data

We removed samples with fewer than 2,500 total reads classified as fungal at the class level (8 samples in the maize dataset, 0 samples in the sorghum and soybean datasets). We further retained only those fungal taxa supported by at least 25 RNA-seq reads in at least one third of all samples within a given dataset, leaving 164, 704, and 246 fungal taxa in the maize, sorghum, and soybean datasets, respectively, including partially redundant taxa classified at multiple taxonomic ranks (Supplementary Table S1; Supplementary Datasets S1-S3). These raw relative abundance values were used for initial community composition analyses, with further normalization applied for association analyses.

Because replicate structure and data availability differed across datasets, genotype-level representation was defined separately for each species. For sorghum, principal components were calculated from log-transformed gene expression (TPM) data. For genotypes with multiple samples, the replicate closest to the genotype-specific centroid in principal component space (PC1–PC5) was retained as the representative. This resulted in a final set of 736 samples representing 736 unique genotypes with both expression and variant data. For maize, replicate samples were removed along with B73HTRHM, a maize genotype closely related to B73, and three genotypes (DK84QAB1, HP72-11, and PHT69) absent from the WiDiv resequencing panel, yielding a final set of 688 samples representing 688 unique genotypes. For soybean, abundance values were averaged across the two genotypes represented by duplicate samples, yielding a final set of 620 samples representing 620 unique genotypes used for transcriptome-wide association studies.

For each fungal taxon, relative abundance values were winsorized at the 1st and 95th percentiles prior to association analysis to reduce the influence of high-magnitude outliers. Because sequencing depth was not optimized for fungal detection, zero abundances in samples passing the minimum read threshold may more plausibly reflect detection limits than confirmed biological absence; as such, zero abundance samples were therefore treated as missing data in all association analyses. Results were not materially affected when analyses were repeated with zero values retained.

### Genome-wide, transcriptome-wide, and expression quantitative trait loci mapping (GWAS, TWAS, and eQTL)

All genome-wide association studies were conducted using the mixed linear model (MLM) as implemented in the rMVP R package (version 1.4.0) [33,34]. Sorghum GWAS used the RNA-seq-derived marker dataset described above, while maize GWAS used a published whole-genome resequencing marker set thinned to 1 SNP per 1 kb using VCFtools (version 0.1.16) [35] to reduce redundancy among highly correlated markers. For each dataset, markers were excluded where fewer than 5% of individuals were homozygous for the minor allele or more than 10% were heterozygous, and the effective number of independent markers (Meff) was estimated using GEC (version 0.266) [36] (Supplementary Table S2). A kinship matrix and three principal components from the respective genetic marker dataset were included as covariates in all GWAS models. Statistical significance was determined using a Bonferroni correction based on Meff, such that markers were considered significant at p < (0.05 / Meff), resulting in species-specific significance thresholds (Supplementary Table S2). Peaks were defined as groups of three or more significant markers within a one-megabase window, with peak density calculated using a sliding window approach (window size 1 Mb, step size 100 bp).

Gene expression was estimated using Kallisto (version 0.46.0) [37] against an index generated from the B73 RefGen_v5 (maize), Wm82.a2.v1 (soybean), or BTx623 v5.1 (sorghum) primaryTranscriptOnly transcript FASTA files obtained from Phytozome [38]. Pseudoalignment outputs were generated as both TPM and raw counts. Raw counts were filtered to remove genes with zero counts in more than 50% of samples, TMM-normalized to counts per million (CPM) using edgeR (version 4.6.3) [39], and log₁₀-transformed prior to use.

Prior to TWAS, transcripts not expressed at ≥0.1 TPM in at least half of samples were excluded. Remaining transcript abundances were rescaled using the approach of Li et al. [40] in which the lowest 5% of samples per gene were assigned a score of 0, the highest 5% a score of 2, and the remaining 90% linearly rescaled between 0 and 2. Associations between rescaled transcript abundances and fungal relative abundances were tested using the compressed mixed linear model (cMLM) in GAPIT (version 3.1) [41,42], with the first three principal components of expression variation included as covariates. Associations were considered significant at a Benjamini-Hochberg false discovery rate [43] of 0.05, applied post-hoc across all transcript-taxon pairs tested.

eQTL analysis was performed for TWAS-significant genes in maize and sorghum (181 and 465 genes, respectively) using log₁₀ CPM TMM-normalized expression values and the same model parameters, marker datasets, and peak-calling criteria as the fungal GWAS. Co-localization between GWAS and eQTL peaks was assessed by permutation test, randomly shuffling GWAS peak positions across the genome 10,000 times and recording overlaps with eQTL peaks within 250 kb. Fold enrichment was calculated as observed overlaps divided by the mean number of permuted overlaps.

### Heritability and variance partitioning

Narrow-sense heritability was estimated using a kinship matrix generated by the calc-kins-direct function in LDAK (version 6.1) [44] with the –reml function and heritability constrained to [0, 1]. Fungal relative abundance values were multiplied by 10,000 prior to analysis to stabilize variance estimation given the inherently small numeric scale of relative abundance data. Only LDAK models that successfully converged were retained for downstream analyses.

### Enrichment analyses

GO and KEGG enrichment analyses were performed using gprofiler2 (version 0.2.4) [45] with all expressed genes in the testing set as the background and FDR correction applied to retain only significant enrichment terms (FDR < 0.05).

### Coexpression and gene regulatory network analyses

Modules of coexpressed genes were identified using WGCNA (version 1.73) [46]. Soft-thresholding powers were selected using the scale-free topology criterion (pickSoftThreshold; powers 1–20). Networks were constructed using a signed topological overlap matrix (TOM) with blockwise module detection (blockwiseModules), applying a minimum module size of 30, a module merging threshold of 0.25, and a maximum block size of 6000 genes to the log₁₀ CPM TMM-normalized expression datasets.

Module eigengenes from the WGCNA-identified modules were correlated with winsorized fungal relative abundances across samples using Pearson correlation with pairwise complete observations. For each correlation, p-values were calculated using the Pearson correlation t-test, with degrees of freedom determined by the number of non-missing observations. P-values were adjusted for multiple testing using the false discovery rate (FDR) method. Modules with significant correlations (FDR < 0.05) were ranked by the median coefficient of determination (R²) across associated fungal taxa, and the top modules were retained for downstream analysis. Functional enrichment results were assigned to each module based on the most significant KEGG pathway enriched among the genes within the module.

Gene regulatory networks were inferred using GENIE3 (version 1.30.0) [47] default random forest parameters (tree-based ensemble method), using the same normalized expression data as input and restricting candidate regulators to transcription factors present in both the expression dataset and the PlantTFDB (soybean) [48] or Grassius (maize, sorghum) [49] transcription factor database for each species.

### Orthology mapping

Orthologous relationships among significant candidate genes were assessed using OrthoFinder (version 3.1.3) [50] with default parameters applied to the B73 RefGen_v5 (maize), Wm82.a2.v1 (soybean), and BTx623 v5.1 (sorghum) primaryTranscriptOnly protein FASTA files obtained from Phytozome.

## Results

### Patterns of fungal relative abundance across crop species

We quantified fungal signal within mRNA-seq datasets generated from mature leaf tissue of 742 maize, 822 sorghum and 622 soybean samples passing quality control, each derived from an independent field study [18–20] (Figure 1A). Among samples passing minimum sequencing thresholds, libraries contained a median of 5,292 (IQR: 4,234–6,711), 59,746 (IQR: 37,003–92,690), and 9,323 (IQR: 7,331–12,501) fungal-assigned reads at the class level for maize, sorghum, and soybean, respectively (Supplementary Figure S1). These results demonstrate consistent recovery of fungal signal across all three crops. Across all datasets, we recovered 29 classes, 66 orders, 131 families, 196 genera and 313 species that met filtering thresholds (≥25 reads in ≥1/3 of samples in at least one dataset; Supplementary Table S1; Supplementary Datasets S1-S3), with patterns of taxonomic overlap varying across datasets and taxonomic ranks (Supplementary Figure S2).

**Figure 1.**
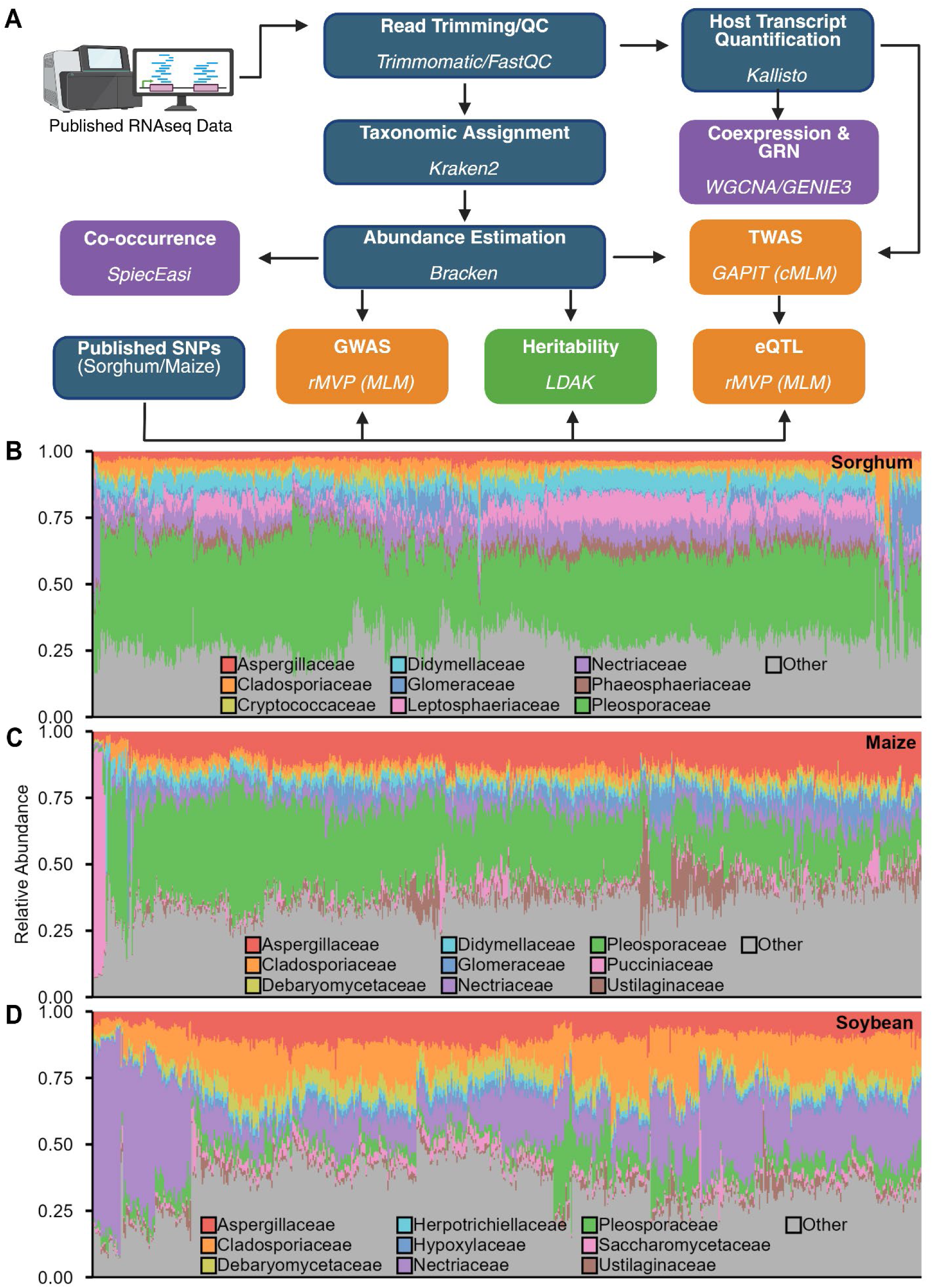
Population scale RNAseq profiling of the leaf mycobiome. **(A)** Schematic illustrating the pipeline used processing and downstream analysis of transcriptomic data. **(B-D)** Variation in observed relative abundance in the nine fungal families with the highest average relative abundance across the samples of each dataset (sorghum, maize and soybean, respectively). Each vertical bar represents one sample and the position of samples on the x-axis was determined by clustering by Bray-Curtis dissimilarity. Schematic illustration created with BioRender.

Despite differences in host species, location and sampling year, several fungal families were consistently abundant, including Aspergillaceae, Cladosporiaceae, Nectriaceae and Pleosporaceae, all of which have also been consistently identified as dominant taxa in crop phyllosphere communities across ITS and metagenomic surveys [51–56], indicating strong concordance between RNA-seq–derived and DNA-based community profiles. Aspergillaceae, Cladosporiaceae, Nectriaceae and Pleosporaceae ranked among the nine most abundant families in all three datasets, while Didymellaceae and Glomeraceae were shared between maize and sorghum, and Debaryomycetaceae and Ustilaginaceae were prominent in maize and soybean (Figure 1B–D). In contrast to these shared dominant taxa, community composition varied substantially among samples within each species, as reflected by clustering based on Bray–Curtis dissimilarity (Figure 1B–D), indicating that fungal relative abundance varies continuously across samples and host genotypes, consistent with quantitative trait variation, which we evaluate explicitly in subsequent analyses. Together, these patterns demonstrate that fungal signal recovered from mRNA-seq data recapitulates key ecological and taxonomic features observed in amplicon- and metagenomics-based studies, supporting its use as a scalable proxy for profiling the leaf mycobiome. Given the rapid expansion of publicly available RNA-seq datasets, this approach enables the reuse of existing transcriptomic data to study plant–microbiome interactions across diverse species, environments, and experimental designs, representing a substantial and previously underutilized resource for large-scale microbiome analyses.

The scale of these datasets enabled construction of co-occurrence networks with greater sample sizes than typically available for phyllosphere studies. Family-level networks revealed partially conserved patterns of connectivity across species, alongside clear host-specific differences in network structure (Figure 2A–C). The total number of significant edges varied across datasets, in part reflecting differences in the number of taxa retained after filtering (Supplementary Table S1).

**Figure 2.**
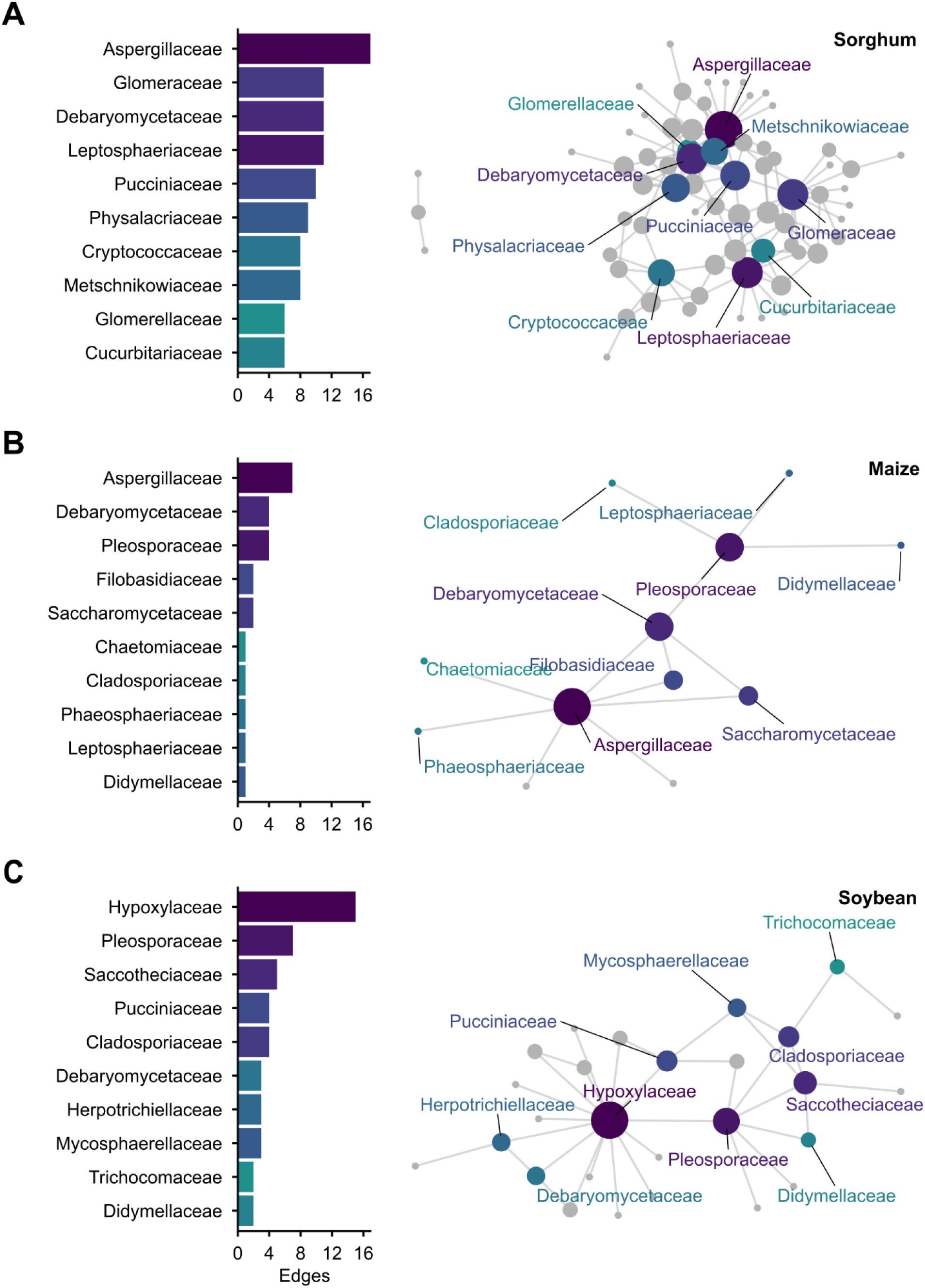
Fungal co-occurrence patterns in the leaf mycobiome. **(A-C)** Fungal co-occurrence networks for sorghum, maize, and soybean, respectively. Nodes represent fungal families, and edges represent significant co-occurrence relationships inferred from network analysis. The ten families with the highest degree (number of edges) in each dataset are labeled. Node size and color both scale with degree centrality within each environment-specific network; values are not directly comparable across panels due to independent network inference and scaling within each dataset.

Several families consistently exhibited high connectivity. Aspergillaceae and Debaryomycetaceae were among the most highly connected taxa in both maize and sorghum, but not in soybean, despite being among the most abundant families in all three datasets (Figures 1B–D, 2A–C). In contrast, Hypoxylaceae displayed the highest connectivity in soybean but was not among the top ten most connected families in maize or sorghum. These results indicate that highly abundant taxa do not necessarily correspond to highly connected taxa and that network structure varies across host species.

### Fungal abundance associates with host gene expression

We performed transcriptome-wide association studies (TWAS) to identify host genes linked to the abundance of individual fungal taxa in maize, sorghum, and soybean (Figure 3; Supplementary Table S4). Across all species, we observed both positive and negative associations between gene expression and fungal abundance, with hundreds to thousands of significant gene-taxon associations: 438 in maize (181 genes × 91 taxa), 4553 in sorghum (465 genes × 486 taxa), and 1692 in soybean (421 genes × 154 taxa). Enrichment analyses revealed that TWAS-significant genes are involved in photosynthesis and energy metabolism, host defense and stress responses, as well as lipid- and fatty acid-related processes likely contributing to cuticular wax formation and plant barrier functions. (Supplemental Figures S3-S6).

**Figure 3.**
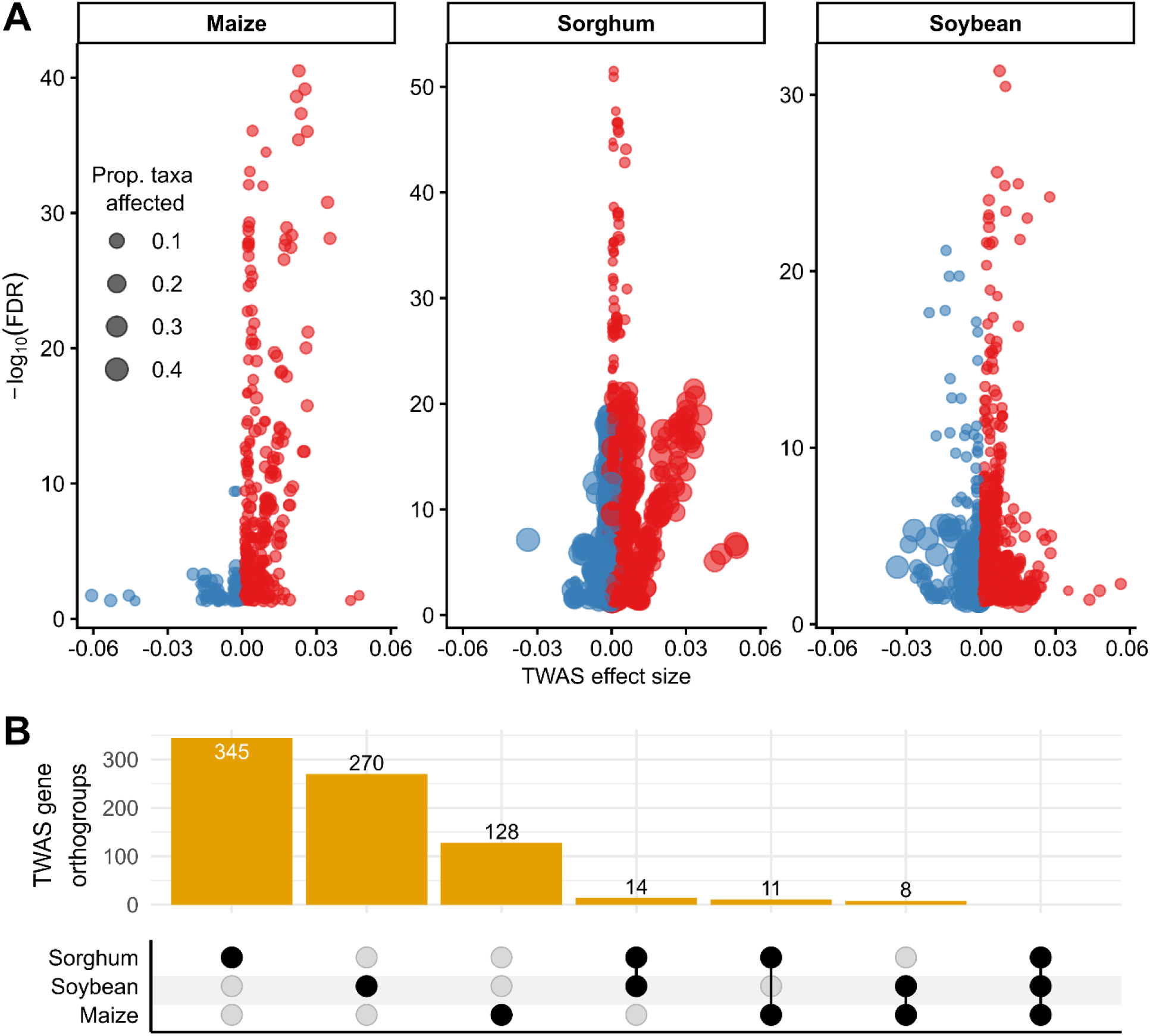
Transcriptome-wide associations identify fungal-linked host genes. **(A)** Volcano plot of TWAS significant gene-fungi associations for each host species. Points indicate a unique gene-fungal association, the size indicates what proportion of fungi within the dataset that individual gene was associated with. Genes with a positive effect are shown in red and negative effects in blue. **(B)** Upset plot comparison of TWAS significant gene orthogroups assignments based on Orthofinder results comparing maize, sorghum and soybean. Intersections indicate number of orthologus genes significant for fungal abundance via TWAS across those species.

The breadth of fungal taxa influenced per gene differed markedly across hosts (Figure 3A). Maize exhibited a distributed genetic architecture, with most genes affecting few taxa (median = 2, mean = 2.42, Gini = 0.38) and a limited number of broad-effect genes (maximum 12 taxa). Sorghum displayed a skewed distribution, with several genes affecting many taxa (median = 2, mean = 9.79, maximum = 300, Gini = 0.77), consistent with a subset of multi-taxa genes driving many associations. Soybean showed an intermediate pattern (median = 3, mean = 4.02, maximum = 94, Gini = 0.42). The top 5% of genes accounted for 15% of associations in maize, 59% in sorghum, and 25% in soybean. Kruskal–Wallis tests confirmed highly significant species differences in taxonomic breadth per gene (χ² = 284.12, df = 2, p < 2.2 × 10⁻¹⁶).

We next quantified the magnitude of gene effects using ΔR², the increase in variance explained by including each gene. Maize genes had the largest effect sizes (mean ΔR² = 0.0623, median = 0.0390), followed by sorghum (mean = 0.0383, median = 0.0294) and soybean (mean = 0.0336, median = 0.0283), indicating that, despite affecting fewer taxa, maize genes exerted stronger individual effects. In maize, effect size and ΔR² were positively correlated with the number of taxa affected (Spearman ρ = 0.368 and 0.252), suggesting that genes with larger effects influence a broader fraction of the fungal community. In contrast, sorghum and soybean showed negative correlations (ρ = −0.315 and −0.386 for effect size), consistent with a pattern in which broad-effect genes are relatively rare. Across all species, ΔR² strongly correlated with statistical significance (ρ = 0.619–0.901), confirming that variance explained predicts robustness of detection.

Finally, comparison of orthologous genes across hosts revealed limited conservation (Figure 3B). Only 14 genes were shared between sorghum and soybean, 11 between maize and sorghum, and 8 between maize and soybean, with no orthogroups detected across all three species (Supplementary Dataset S4). These findings suggest that, while some host genetic mechanisms influencing fungal abundance may be conserved, the majority of associations are species-specific, reflecting divergent host–microbiome interactions.

### Host gene modules predict fungal associations

To examine how host transcriptional programs relate to leaf-associated fungi, we constructed WGCNA coexpression networks for maize, sorghum, and soybean, grouping genes into modules of tightly coexpressed loci. This allowed us to quantify module-level associations with fungal abundance. Across species, median module–fungal correlations were low (Figure 4A; median R²: maize 0.0069, sorghum 0.019, soybean 0.034), indicating that most modules explain only a small fraction of fungal variation. Nonetheless, specific module–fungus pairs in sorghum and soybean reached strong correlations (R² > 0.4), despite low median module R², reflecting concentrated effects of a few modules rather than broad influence across all module members.

**Figure 4.**
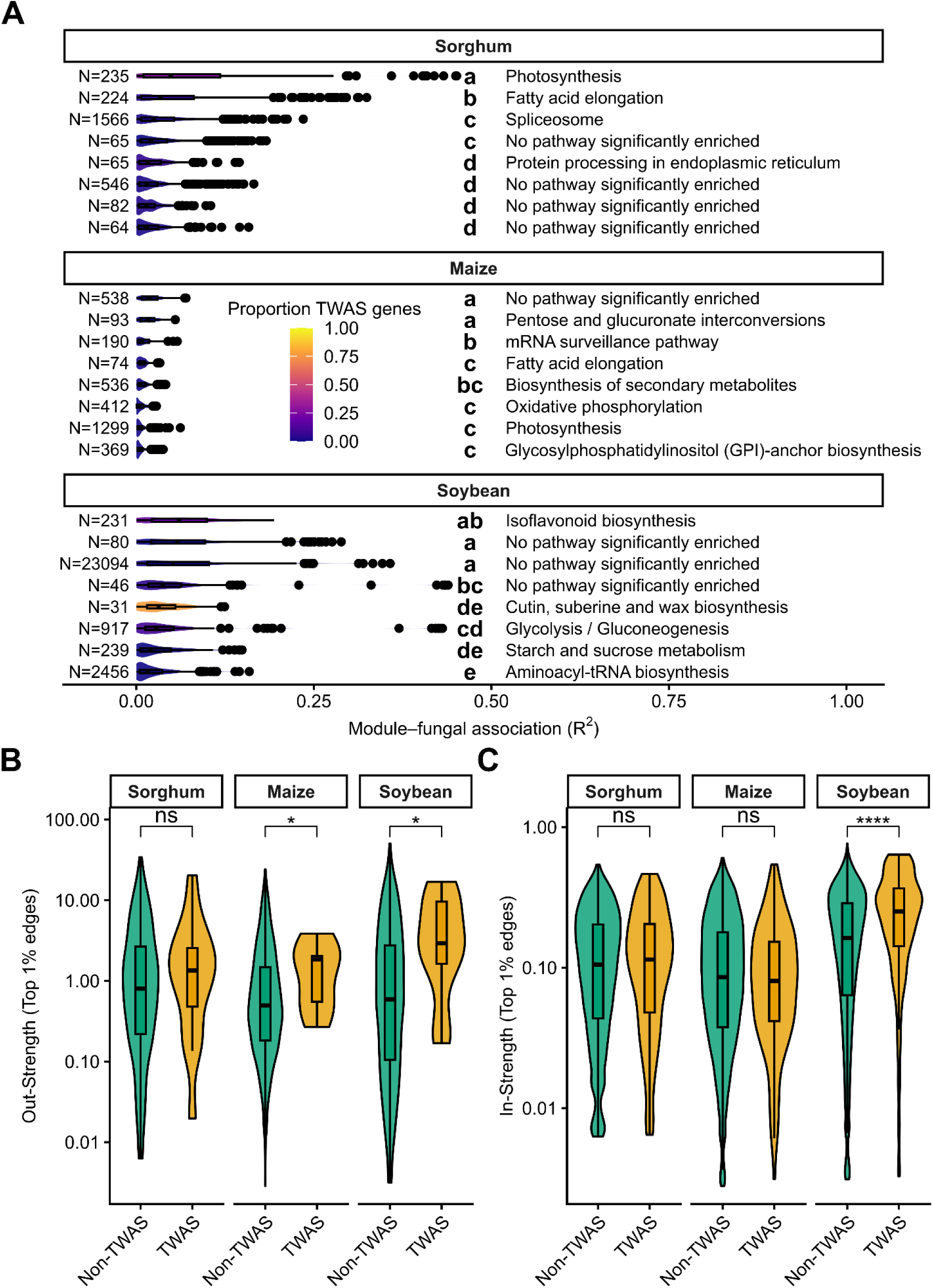
Host transcriptional architecture underlying fungal association. **(A**) Distributions of R^2^ values of Pearson correlation between fungal abundance and WGCNA module eigengene values for top 8 modules for each species by median R^2^. Letters indicate significance via one-way ANOVA with Tukey’s HSD. Annotations on the right indicate the most significantly enriched KEGG term for genes within each module. N values represent number of genes within each module. Color of violins indicates proportion of genes within each module which were significant for at least one fungal taxon via TWAS. **(B)** Comparison in GENIE3 network out-strength distributions across three host species for genes either significantly or not significantly associated with any fungal taxa via TWAS (*p<0.05 via Wilcoxon test; NS = no significant difference). **(C)** Comparison in GENIE3 network in-strength distributions across three host species for genes either significantly or not significantly associated with any fungal taxa via TWAS (****p<0.0001 via Wilcoxon test; NS = no significant difference).

Sorghum and soybean modules showed higher proportions of fungi with significant correlations (median proportion significant 0.415 and 0.461, respectively; Figure 4A, violin shading indicates TWAS enrichment). Top associations in sorghum occurred in modules enriched for photosynthesis and fatty acid metabolism, while soybean modules exhibited strong correlations across diverse metabolic pathways. By contrast, maize modules displayed fewer (∼11%) significant module–fungus associations, consistent with a more distributed or polygenic influence on fungal variation.

Across species, module enrichment for TWAS-identified genes was weakly correlated with module–fungal R² and did not reach statistical significance (Spearman ρ = −0.38–0.36, p > 0.35; Figure 4A, violin shading), suggesting that the presence of TWAS genes alone does not predict a module’s influence. Network centrality analysis with GENIE3 further indicated that TWAS genes generally exhibit lower outgoing regulatory strength but slightly higher incoming strength relative to non-TWAS genes (Figure 4B–C), consistent with a role as regulatory targets rather than dominant hubs. Wilcoxon tests confirmed significant differences in out-strength for maize and soybean (p < 0.05), whereas sorghum showed similar trends without statistical significance.

These results reveal that fungal associations in maize are broadly distributed across the transcriptome, whereas sorghum and soybean display more localized effects driven by specific coexpressed modules. This contrast in module-level association strength and network centrality underscores species-specific differences in the transcriptional architecture underlying fungal community variation.

### Genomic architecture of mycobiome regulation differs in maize and sorghum

To quantify host genetic contributions to leaf-associated fungal communities, we performed GWAS and eQTL mapping of TWAS-significant genes, restricting analyses to maize and sorghum due to the lack of high-density genotype data for soybean. Narrow-sense heritability (h²) of fungal relative abundance spanned a broad range in both species (Figure 5A; Supplementary Table S3), with a higher central tendency in maize (median h² = 0.31, IQR 0.20–0.40) than in sorghum (median h² = 0.14, IQR 0.09–0.23), a pattern also reflected in the full distribution (5–95% range: 0.09–0.59 vs. 0.03–0.47). Nearly all taxa exhibited non-zero heritability (100% in maize; 98.0% in sorghum), and a greater proportion exceeded h² > 0.1 in maize (94.4% vs. 68.4%).

**Figure 5.**
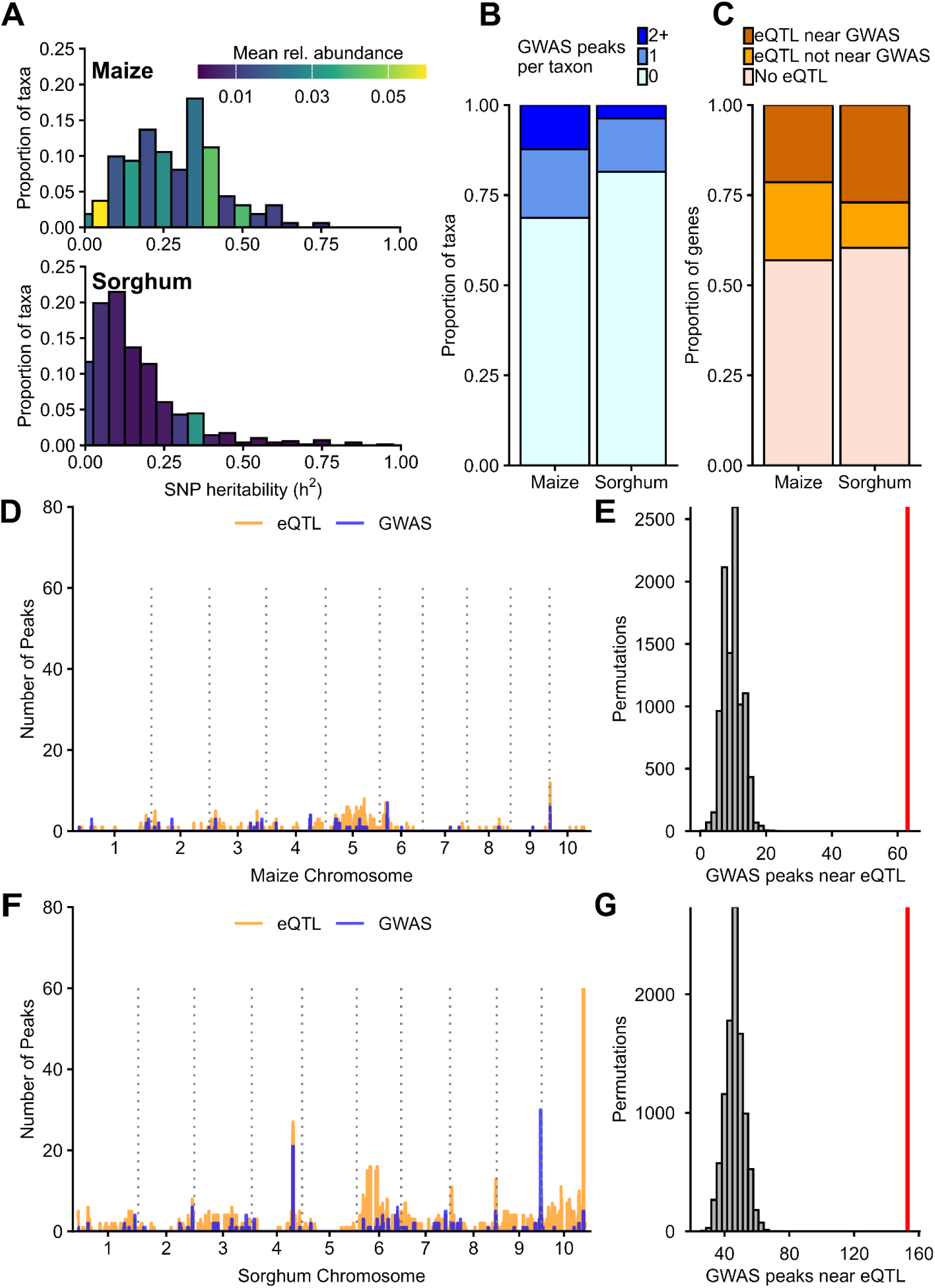
Genome-wide distribution of GWAS and eQTL peaks in maize and sorghum. **(A)** Distributions of narrow-sense heritability (h²) for relative abundance of fungal taxa in maize (top) and sorghum (bottom). Taxa were binned in 0.05 h² intervals, and bar height indicates the proportion of taxa within each bin. Bar color denotes the mean relative abundance of all taxa in that bin. **(B)** Proportion of fungal taxa with 0, 1, or ≥2 significant GWAS peaks per taxon in maize and sorghum. **(C)** Proportion of TWAS significant genes with expression quantitative trait loci (eQTL) that colocalize with GWAS peaks (within ±250 kb), eQTL not near GWAS peaks, or no detected eQTL in maize and sorghum. **(D)** Maize sliding-window plots showing the number of GWAS (blue) and eQTL (orange) peaks across chromosomes, where eQTL peaks are derived from TWAS-significant genes. Vertical dotted lines indicate chromosome boundaries. **(E)** Histograms showing the overlap of GWAS peaks randomly placed across the genome (10,000 permutations), with the locations of the TWAS-eQTL peaks. True observed number of overlaps with eQTL peaks indicated by red line. **(F)** Sorghum sliding-window plots for GWAS and TWAS-eQTL peaks and **(G)** corresponding permutation histograms.

Heritability was not meaningfully correlated with mean relative abundance in maize (Spearman ρ = −0.078, p = 0.325) and showed at most a very weak relationship in sorghum (ρ = 0.080, p = 0.046), indicating that host genetic control is not strongly structured by taxon abundance in either species. Across taxa shared between species, continuous heritability estimates were not significantly correlated (Spearman ρ = −0.15, p = 0.07). Using a predefined threshold for detectability (h² > 0.1), taxa classified as heritable in one species were not more likely to be classified as heritable in the other (Fisher’s exact test, p = 0.45), indicating no evidence for conserved classification of detectable heritability at the level of individual fungal taxa between maize and sorghum.

Despite the more distributed transcriptional architecture observed in maize (Figure 4A), GWAS of fungal relative abundance revealed more pervasive genetic control. Consistent with this detectability framework, taxa with higher heritability were significantly more likely to yield GWAS associations (logistic regression, p = 0.0077), indicating that mapping success reflects biologically meaningful host genetic control rather than stochastic detection. In absolute terms, more taxa exhibited significant associations in sorghum (130 taxa; 179 total peaks) than in maize (51 taxa; 87 total peaks), consistent with higher overall taxonomic richness (Supplementary Table S1). However, when normalized by the number of taxa tested, a larger proportion of taxa in maize exhibited genetic associations, with fewer taxa lacking significant associations (68.7% vs. 81.5%) and more taxa exhibiting one (19.0% vs. 14.8%) or multiple associated loci (12.3% vs. 3.7%) (Figure 5B). The number of associations per taxon was also higher in maize (mean 0.53 vs. 0.26), with more extreme cases (up to 10 peaks per taxon in maize vs. 8 in sorghum), supporting a more polygenic architecture. All significant loci are reported in Supplementary Table S5.

In contrast, sorghum exhibited a more concentrated genomic architecture. Sliding-window analyses revealed regions of elevated peak density in both species (Figure 5D,F), with higher raw peak counts per window in sorghum (GWAS: mean 0.24, max 30; eQTL: mean 1.10, max 76) than in maize (GWAS: mean 0.041, max 7; eQTL: mean 0.20, max 12), although these differences are influenced by variation in genome size and total peak number. To enable direct comparison, we quantified the distribution of peaks across genomic bins using normalized metrics. Sorghum displayed a more uneven distribution of peaks than maize, with higher Gini coefficients for both GWAS (0.46 vs. 0.31) and eQTL peaks (0.46 vs. 0.30), indicating stronger concentration of associations within a subset of genomic regions. Consistent with this, the top 10% of genomic windows accounted for a larger fraction of peaks in sorghum (∼39–41%) than in maize (∼25–27%), highlighting the concentrated nature of sorghum fungal regulatory hotspots.

Given the contrasting GWAS architectures, we next examined whether regulatory variation showed similar species-level structure. We assessed the regulatory basis of TWAS signals by testing for colocalization between fungal abundance GWAS peaks and eQTLs of TWAS-identified genes (Supplementary Table S6). A minority of TWAS-significant genes colocalized with GWAS peaks in both species, though this proportion was modestly higher in sorghum (27.0%) than in maize (21.4%), while a comparable fraction showed eQTLs distal to GWAS loci (12.6% vs. 21.7%). A substantial proportion of TWAS genes lacked detectable eQTLs (60.4% in sorghum; 56.9% in maize) (Figure 5C). However, permutation tests revealed that overlap between GWAS peaks and eQTLs was significantly greater than expected by chance in both species (maize: 6.39-fold enrichment; sorghum: 3.3-fold enrichment; both p < 1×10⁻⁴), supporting a model in which TWAS associations are primarily driven by cis-regulatory variation rather than downstream transcriptional responses.

Together, these results demonstrate that host genetic control of the leaf mycobiome is widespread and substantial in both maize and sorghum but arises from distinct architectures. Maize exhibits a distributed, polygenic architecture characterized by many loci of modest effect, whereas sorghum shows a more modular architecture with concentrated, high-density genomic regions influencing multiple taxa.

### Broad-spectrum fungal regulatory loci are driven by cis-regulatory variation in membrane-bound receptor genes

Among the fungal regulatory hotspots identified in sorghum, we prioritized a locus on chromosome 4 (Chr04: 57.99-59.96 Mb) due to its strong multi-trait signal. This region was associated with 21 fungal taxa via GWAS and overlapped an interval (Chr04: 58.12-59.56 Mb) containing eQTL signals for 27 genes, all of which were also significant in TWAS (Supplementary Table S7). Among the associated taxa was *Clavicipitaceae*, a fungal family containing several ecologically and agronomically relevant members including *Metarhizium* spp., *Claviceps* spp., and *Epichloë* spp. [57–59].

Within this interval, a leucine-rich repeat receptor-like kinase (LRR-RLK; *Sobic.004G214900*) emerged as the primary candidate gene. This gene was significant in TWAS for *Clavicipitaceae* and 13 additional fungal taxa (Supplemental Table S8), co-localized with the GWAS peak, and harbored a strong cis-eQTL (Figure 6A–C). Genotype at the peak GWAS SNP (Chr04_58988275) was strongly associated with both *Sobic.004G214900* expression (median 0.50 vs. 1,584 TPM; *P* < 2.2 × 10⁻¹⁶, Wilcoxon test; Figure 6D) and fungal relative abundance (median 0.0024 vs. 0.0241; *P* < 2.2 × 10⁻¹⁶; Figure 6E). Expression of *Sobic.004G214900* was also positively correlated with *Clavicipitaceae* abundance (Spearman ρ = 0.196, *P* = 9.49 × 10⁻⁸), consistent with a genotype-mediated regulatory relationship.

**Figure 6.**
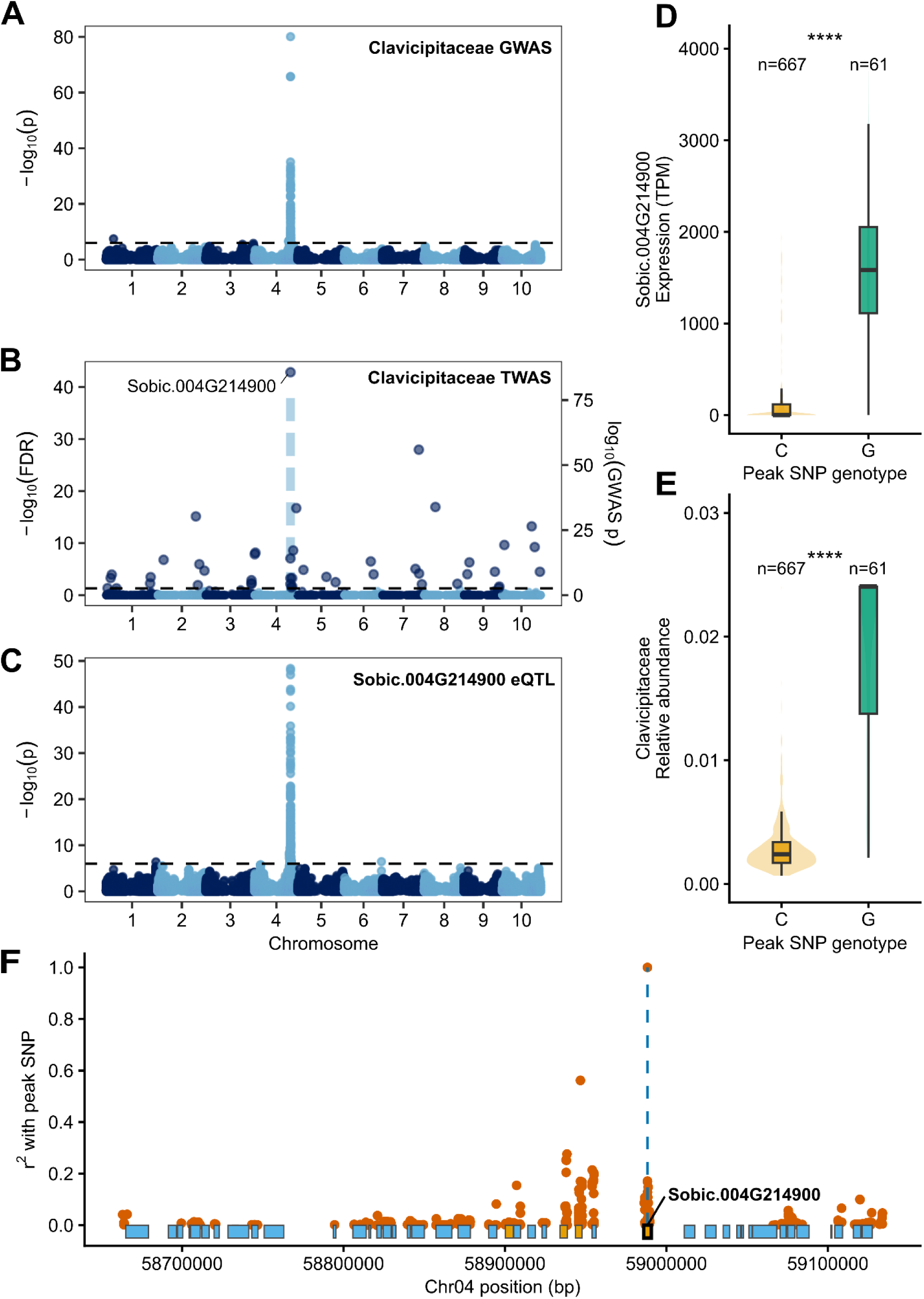
Integrative genetic and genomic analysis of a Chr04 locus associated with fungal relative abundance and gene expression. **(A)** Manhattan plot of the genome-wide association study (GWAS) for Clavicipitaceae relative abundance, showing genomic positions of SNPs across all chromosomes (x-axis) and the −log10(p-value) of association (y-axis). The horizontal dashed line indicates the Bonferroni-adjusted genome-wide significance threshold (0.05/50,150). **(B)** Manhattan plot of the transcriptome-wide association study (TWAS) for Clavicipitaceae relative abundance. The horizontal dashed line indicates the significance threshold (FDR = 0.05). **(C)** Manhattan plot of the expression quantitative trait locus (eQTL) analysis for Sobic.004G214900, showing SNP associations with expression levels of this gene across the genome. The horizontal dashed line indicates the Bonferroni-adjusted genome-wide significance threshold (0.05/50,150). **(D)** Boxplots of normalized expression levels of Sobic.004G214900 stratified by genotype (homozygous individuals only) at the peak SNP identified in the Clavicipitaceae relative abundance GWAS (Chr04_58988275). Stars indicate P < 2.2 × 10⁻¹⁶ (Wilcoxon test). **(E)** Boxplots of Clavicipitaceae relative abundance stratified by genotype (homozygous individuals only) at the peak SNP identified in the Clavicipitaceae relative abundance GWAS (Chr04_58988275). Stars indicate P < 2.2 × 10⁻¹⁶ (Wilcoxon test). **(F)** Linkage disequilibrium (LD) landscape surrounding the chromosome 4 peak SNP. Points represent pairwise LD (r²) between each SNP in the region and the peak SNP. The vertical dashed line marks the position of the GWAS peak SNP (Chr04_58988275). Rectangles below the axis denote annotated genes within the interval (BTx623 V5 reference genome), with orange indicating TWAS significance; the focal gene Sobic.004G214900 is highlighted.

Linkage disequilibrium analysis indicated that this gene resides within a low-LD interval containing few other TWAS-significant genes, supporting it as the primary candidate underlying the association (Figure 6F).

Additional loci with partially similar signatures were identified but showed less clear candidate resolution. A region on chromosome 9 (Chr09: 61.62-63.28 Mb) was associated with 30 taxa (Supplementary Table S9), with nearby genes including a pentatricopeptide repeat (PPR) protein (*Sobic.009G254200*) and another LRR-containing protein (*Sobic.009G254400*), although the latter exhibited minimal expression across samples. Another region on chromosome 10 (Chr10: 58.88-60.74 Mb) was associated with the expression of 77 TWAS-significant genes (Supplementary Table S10) but lacked a clearly resolved candidate gene.

Together, these results identify discrete loci in sorghum where cis-regulatory variation in host genes drives coordinated changes in gene expression and fungal colonization, with the strongest-supported example implicating a membrane-bound receptor kinase as a key regulator of fungal abundance. These findings support a model in which variation in host receptor-mediated signaling may influence permissiveness to fungal colonization, producing large, taxa-specific effects at individual loci. This architecture contrasts with the more diffuse, polygenic control observed in maize, highlighting species-specific differences in the genetic regulation of plant-associated fungal communities. More broadly, these results demonstrate that host genetic control of microbiome composition can be resolved through integrated analysis of transcriptomic and association data, establishing a scalable framework for leveraging existing RNA-seq datasets to dissect plant–microbe interactions across diverse systems.

## Discussion

Despite growing recognition that host plant genetics shapes the composition of associated microbial communities, efforts to identify specific genomic loci controlling microbial abundance have met with limited success. Studies of leaf-associated bacteria and fungi in maize, sorghum, and Arabidopsis using amplicon-based profiling have recovered few significant associations, with most traits exhibiting low heritability and or with no signals surviving stringent correction for multiple testing [5,6,9,11]. Here, leveraging population-scale RNA-seq datasets not originally generated for mycobiome profiling, we identify hundreds of significant host genetic and transcriptomic associations with leaf fungal abundance across three crop species, far exceeding prior efforts and suggesting that host genetic control of the phyllosphere mycobiome has been systematically underestimated. Although paired ITS and RNA-seq data from identical field-collected tissues are not available at population scale, methodological validity is supported by concordance of dominant community members with established amplicon and metagenomic surveys [51–56], recovery of known pathogen recognition receptors with directionally concordant mutant phenotypes, and significant enrichment of GWAS-eQTL colocalization independent of community profiling assumptions.

Differences in sequencing depth alone are unlikely to explain this improvement. While fungal read counts were relatively high in sorghum (median 59,746 fungal reads per sample), maize read depths (median 5,292 reads per sample) were comparable to or below thresholds used in several amplicon studies [5,7,9] yet yielded robust associations. Instead, we argue that two features of RNA-seq–based profiling jointly enhance detection power. First, by capturing only polyadenylated transcripts, this approach enriched for fungal transcripts from metabolically active fungi, avoiding the signal dilution introduced by dormant spores and residual environmental DNA that confound DNA-based methods [15,17]. Second, the lower PCR amplification in RNA-seq library preparation (∼8–10 cycles versus 25–35 for amplicon workflows) reduces stochastic noise, allowing read counts to more faithfully reflect true biological abundance. The resulting improvement in quantitative accuracy likely underlies both the stronger individual associations and the higher heritability estimates we observe relative to previous studies. Consistent with this interpretation, a prior study of metabolically active maize leaf bacteria using RNA-based amplicons reported few significant associations and low heritability for individual taxa [9], suggesting that RNA as a substrate alone is insufficient and that the reduced amplification inherent to shotgun library preparation is a key contributing factor.

The population sizes employed here (688 to 736 individuals per species) also substantially exceed those of prior microbiome GWAS efforts, which have typically ranged from 175 to 323 genotypes [5–7,9,11,12]. While power contributes to discovery, the distribution of effect sizes we observe, including several loci individually explaining 20–40% of variance in fungal abundance, suggests that some associations would likely have been detectable at smaller sample sizes had quantification been sufficiently accurate. Together, these factors point to measurement precision as a primary limiting factor in previous microbiome GWAS, with implications for how future studies should be designed.

The genetic architectures we observe in maize and sorghum are distinct despite broadly similar fungal community composition. Maize exhibits a distributed, polygenic architecture in which many loci of modest effect collectively shape fungal abundance across taxa. Sorghum, by contrast, displays a concentrated architecture dominated by a small number of genomic hotspots, each associated with the abundance of many fungal taxa simultaneously, mirroring, at the phyllosphere level, a similar pattern previously reported for sorghum rhizosphere bacterial communities using relaxed significance thresholds [5]. Although the maize and sorghum marker sets differ in ascertainment strategy, RNA-seq-derived markers have previously been shown to perform as substitutes for WGS markers in association mapping while enabling retention of more genotypes [18]. The architectural differences we observe between species are independently recapitulated in the transcriptomic data, suggesting they reflect genuine biological differences rather than marker density artifacts. Whether these architectural differences reflects fundamental differences in the genetic basis of immunity and microbiome regulation between these two grasses, differences in pathogen pressure across environments, or other factors remains an open question, but the contrast is robust to normalization for differences in total peak number, demonstrating that the concentration of peaks in sorghum is not simply a consequence of having more total loci (Supplemental Figure S7).

A key interpretive challenge in transcriptome-wide association studies of microbiome traits is causal direction: does host gene expression drive fungal abundance, or does fungal colonization alter host transcription? The integration of GWAS, TWAS, and eQTL mapping presented here provides strong evidence favoring the former interpretation in at least a subset of cases. We find that overlaps between fungal GWAS peaks and eQTLs for TWAS-significant genes are significantly enriched beyond chance in both maize and sorghum (6.4-fold and 3.3-fold, respectively; both p < 1×10⁻⁴), indicating that shared genetic variants simultaneously regulate host transcript abundance and fungal colonization. Under a reverse-causation model, where fungal presence induces host gene expression changes, one would not expect this pattern of cis-regulatory colocalization. These results are consistent of a model in which natural variation in cis-regulatory sequences modulates the expression of host immune, signaling, or metabolic genes, which in turn influence the permissiveness of the leaf environment to specific fungal colonizers. While functional dissection of individual candidate loci through genetic manipulation will be an important next step in establishing causality, the convergence of independent genomic signals at biologically plausible genes provides a strong observational basis for prioritizing specific loci and hypotheses for such follow-up work.

The strongest example of this architecture occurs at a sorghum chromosome 4 locus where convergent GWAS, TWAS, and eQTL signals implicate a leucine-rich repeat receptor-like kinase (*Sobic.004G214900*) as a regulator of abundance across 14 fungal taxa simultaneously, including Clavicipitaceae. LRR-RLKs constitute the largest family of immune receptors in plants and are well established as mediators of both pathogen recognition and microbiome homeostasis[60–63]. The allele associated with higher Clavicipitaceae abundance is also associated with dramatically higher expression of this receptor (median 0.50 vs. 1,584 TPM), suggesting a permissive rather than restrictive role, where higher receptor activity may facilitate rather than suppress colonization by this family, perhaps through signaling pathways that promote endophytic accommodation. This interpretation is consistent with the ecology of Clavicipitaceae, which includes not only pathogens such as *Claviceps* but also beneficial endophytes, including *Epichloë* spp. and the entomopathogenic genus *Metarhizium*, increasingly recognized as a plant-associated endophyte with potential roles in indirect herbivore defense [57]. Notably, this signal is recapitulated at finer taxonomic resolution, with even stronger associations observed for *Metarhizium* at the genus and species level (e.g., *M. album*), indicating that the effect is not driven solely by broad family-level classification. While we cannot resolve whether these reads correspond to insect-pathogenic strains, the association with a host receptor allele raises the possibility that receptor-mediated signaling influences the abundance of entomopathogenic endophytes in field populations, a hypothesis that warrants targeted investigation.

Co-occurrence network analysis revealed partially conserved but species-specific patterns of fungal connectivity across the three crop systems. Notably, several of the most highly connected families, including Aspergillaceae and Hypoxylaceae, contain endophytic taxa that are well-documented producers of secondary metabolites with antifungal activity such as azaphilones and griseofulvin [64–66]. Their position as network hubs is therefore not only statistically robust but biologically interpretable: taxa capable of chemically suppressing or modulating the growth of competitors are precisely those expected to exert disproportionate influence over community structure. This correspondence between known secondary metabolite production and network centrality suggests that the co-occurrence patterns we observe, though inferred from observational data, reflect ecologically meaningful interactions rather than stochastic co-detection alone.

Across all three species, TWAS identified hundreds of genes whose expression predicts fungal abundance, with functional enrichment for energy metabolism, defense response, and lipid- and fatty acid-related processes likely contributing to cuticular wax formation and leaf barrier functions. Consistent with this defense enrichment, among maize TWAS associations we recover *rlk10* (*Zm00001eb293660*), a leucine-rich repeat receptor-like kinase previously characterized as a fungal-induced receptor involved in pathogen recognition [67]. Loss-of-function mutants exhibit enhanced susceptibility to the necrotrophic pathogen *Cochliobolus heterostrophus*, alongside reduced jasmonic acid and phytoalexin accumulation, but increased resistance to the hemibiotrophic pathogen *Fusarium graminearum* [67], indicating that this receptor can have opposing effects depending on pathogen lifestyle. Notably, *rlk10* has also been independently identified as a candidate locus for southern corn rust resistance in GWAS of Chinese maize germplasm [68], and as a hub gene in co-expression networks responsive to multiple pathogen stresses [69], further supporting its central role in biotic stress responses. In our dataset, *rlk10* expression was associated with the abundance of seven fungal taxa with a directionally consistent pattern: positive associations with Puccinia and its encompassing higher-order taxonomic groups (Pucciniaceae, Pucciniales, and Pucciniomycetes), and negative associations with Dothideomycetes and Pleosporales, the class and order containing *Cochliobolus*. This directional concordance between natural expression variation and published loss-of-function phenotypes, recovered from leaf tissue sampled prior to visible disease development, provides independent corroboration that TWAS of RNA-seq–derived fungal abundance captures biologically meaningful host–pathogen relationships and suggests that *rlk10* mediates opposing effects on phylogenetically distinct fungal groups, potentially reflecting differential responses to fungal trophic strategies mediated through divergent immune signaling pathways.

While individual associations like *rlk10* illuminate specific host–pathogen relationships, the limited conservation of TWAS-significant genes across species, with no orthogroups shared across all three, suggests that the specific molecular players underlying plant–fungal interactions have diverged substantially. This pattern aligns with the rapid birth–death evolution typical of plant immune gene families, where functional roles are maintained through repeated, independent recruitment of paralogs rather than conservation of orthologs [63,70]. Practically, this species-specificity implies that the host genetic levers available to modulate mycobiome composition are largely crop-specific, limiting the direct transferability of candidate genes across systems. Nevertheless, the conservation of broad functional categories despite divergence at the gene level, indicates that shared ecological pressures consistently shape plant–fungal interactions across grasses, even as the underlying genetic mechanisms remain evolutionarily labile.

More broadly, this study demonstrates that the hundreds of thousands of publicly available plant RNA-seq datasets, generated primarily for transcriptomic rather than microbiome purposes, represent a largely untapped resource for studying plant-microbe interactions at population scale. The framework we describe here, applying standard taxonomic classification pipelines to existing polyA-enriched libraries, requires no additional experimental investment and scales with the continued growth of public sequence repositories. As population-scale transcriptomic datasets expand across additional crop species and environments, this approach offers a tractable path toward understanding how host genetic diversity shapes microbiome composition across the breadth of agricultural ecosystems, a question that purpose-built microbiome studies, given their cost and scale requirements, are unlikely to address comprehensively in the near term. The strong, biologically interpretable associations we recover across three species and multiple analytical frameworks suggest that the influence of plant genetics on microbiome composition has been substantially underestimated, and that re-analysis of existing transcriptomic resources could rapidly accelerate discovery.

## Data Availability Statement

Previously published RNA sequencing data used for this study are publicly available through the European Nucleotide Archive (ENA) under accession numbers PRJEB83049 (sorghum) and PRJEB67964 (maize), and through the Genome Sequence Archive (GSA, Beijing Institute of Genomics Data Center) under accession CRA009979 (soybean). For maize and sorghum, genetic marker data were obtained from publicly available repositories (Dryad and Figshare, respectively). The prebuilt Kraken2 classifier database is available from the Langmead Lab AWS index repository. All code used for analysis and visualization is available on GitHub.

Links:

- ENA (sorghum): PRJEB83049
- ENA (maize): PRJEB67964
- GSA (soybean): CRA009979
- Dryad dataset: https://doi.org/10.5061/dryad.bnzs7h4f1
- Figshare dataset: https://doi.org/10.6084/m9.figshare.27936195
- Kraken2 database: https://benlangmead.github.io/aws-indexes/k2
- GitHub repository: https://github.com/ccolvinmmiv/RNAseq_phyllosphere_mycobiome

## Supporting information

List of Supplementary Materials

## Acknowledgments

We thank the Penn State Institute for Computational and Data Sciences and the Penn State One Health Microbiome Center for providing computational resources and support for this project. We also thank James Schnable and Jensina Davis (University of Nebraska–Lincoln) for discussions on association analyses, as well as Maria Alejandra Gil Polo and Sergio Pérez-Limón (The Pennsylvania State University) for their feedback on the manuscript.

## Author Contribution Statement

CC and SC designed experiments. CC performed research. CC analyzed data. CC and SC wrote, reviewed and edited the manuscript.

