## Supplementary Figures S1-S7 for "Population-scale transcriptomics reveals host genetic control of phyllosphere fungal communities"

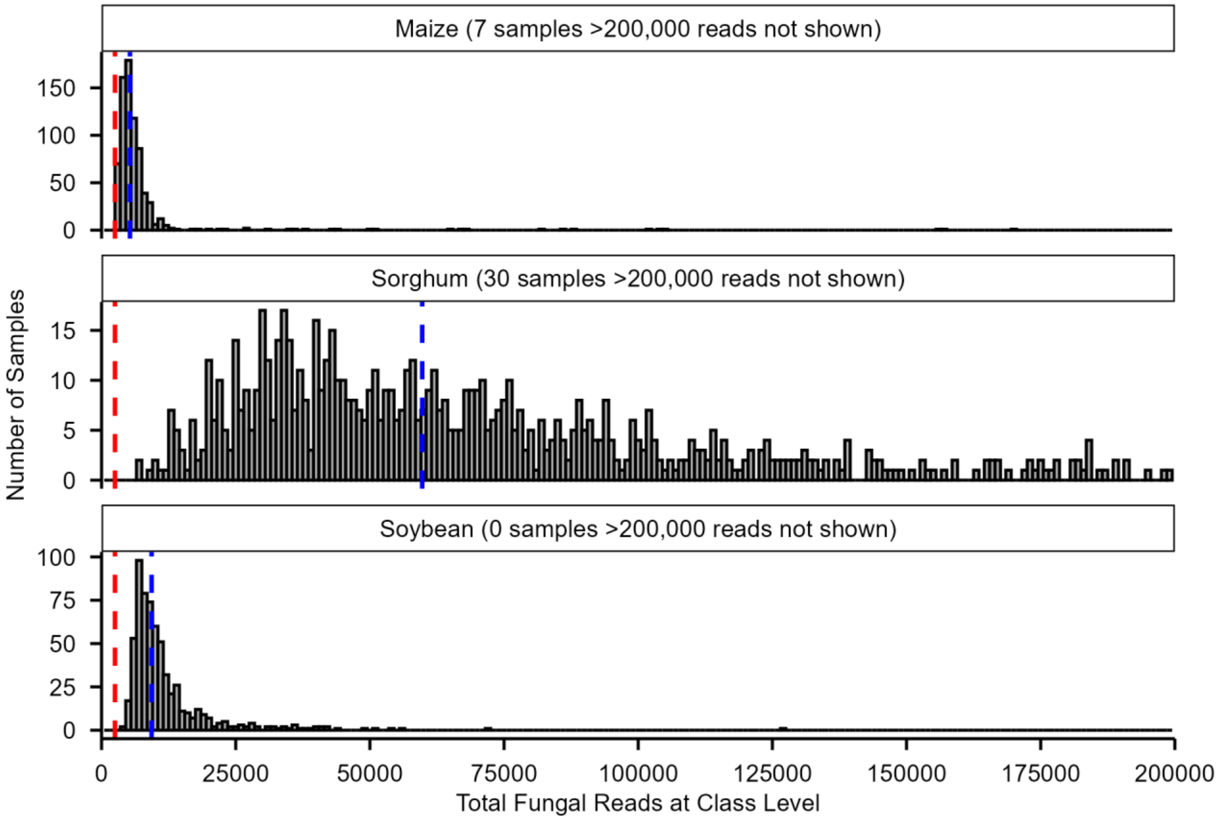


**Supplementary Figure S1. Distribution of fungal read counts per sample across datasets.** Histograms show total fungal reads per sample for each environment (faceted panels). Samples exceeding 200,000 reads are not displayed, with counts of omitted samples indicated in each panel. Dashed vertical lines denote the minimum read threshold (red; 2,500 reads) and the median read count (blue) per dataset.

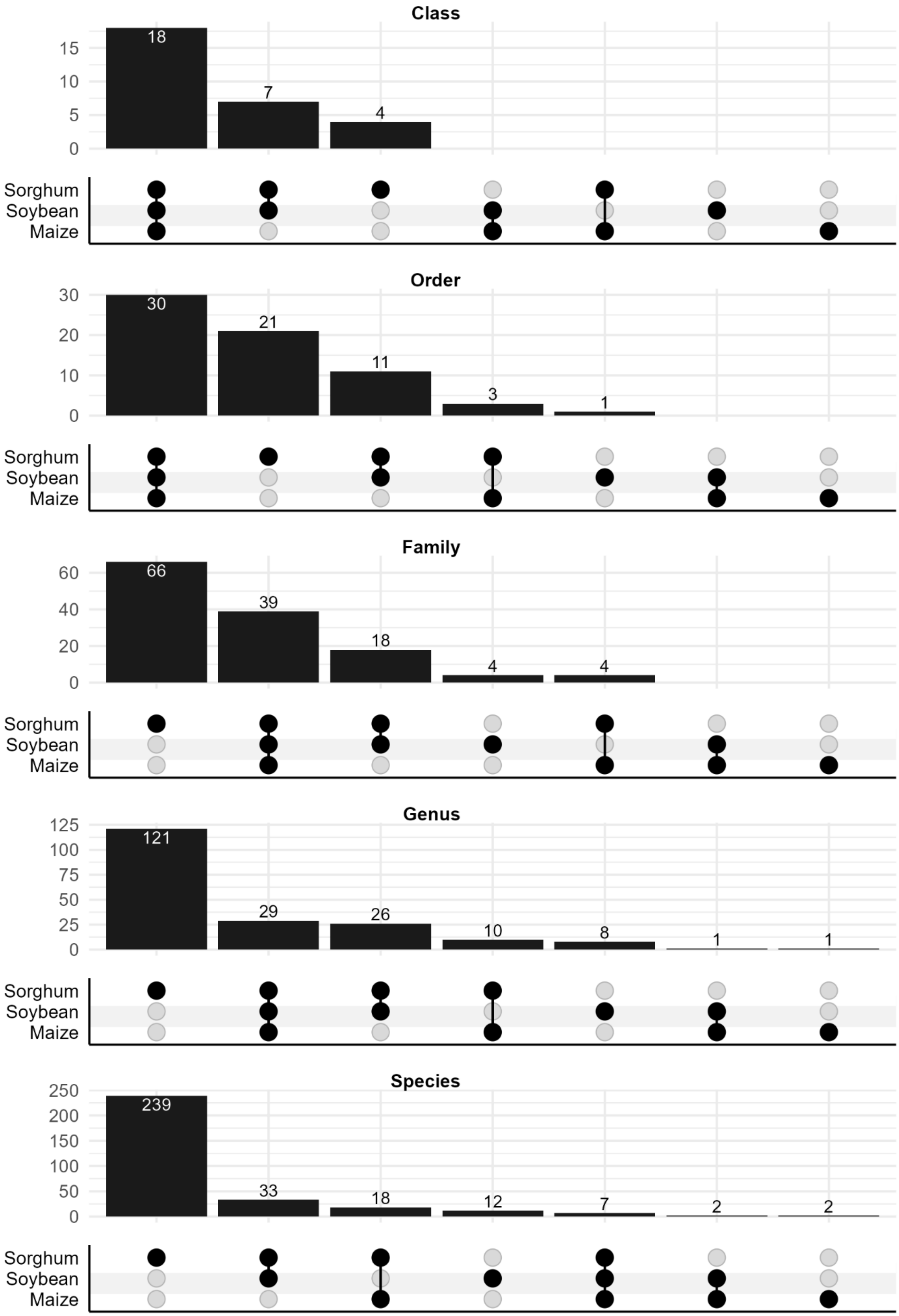

Class
18
7
4
15
10
5
0
Sorghum
Soybean
Maize
Order
30
21
11
3
1
30
20
10
0
Sorghum
Soybean
Maize
Family
66
39
18
4
4
60
40
20
0
Sorghum
Soybean
Maize
Genus
121
29
26
10
8
1
1
125
100
75
50
25
0
Sorghum
Soybean
Maize
Species
239
33
18
12
7
2
2
250
200
150
100
50
0
Sorghum
Soybean
Maize

**Supplementary Figure S2.** Overlap of fungal taxa detected across maize, sorghum, and soybean for each taxonomic level. Plots are shown for Classes, Orders, Families, Genera, and Species. Bars represent the number of taxa present in each combination of hosts.

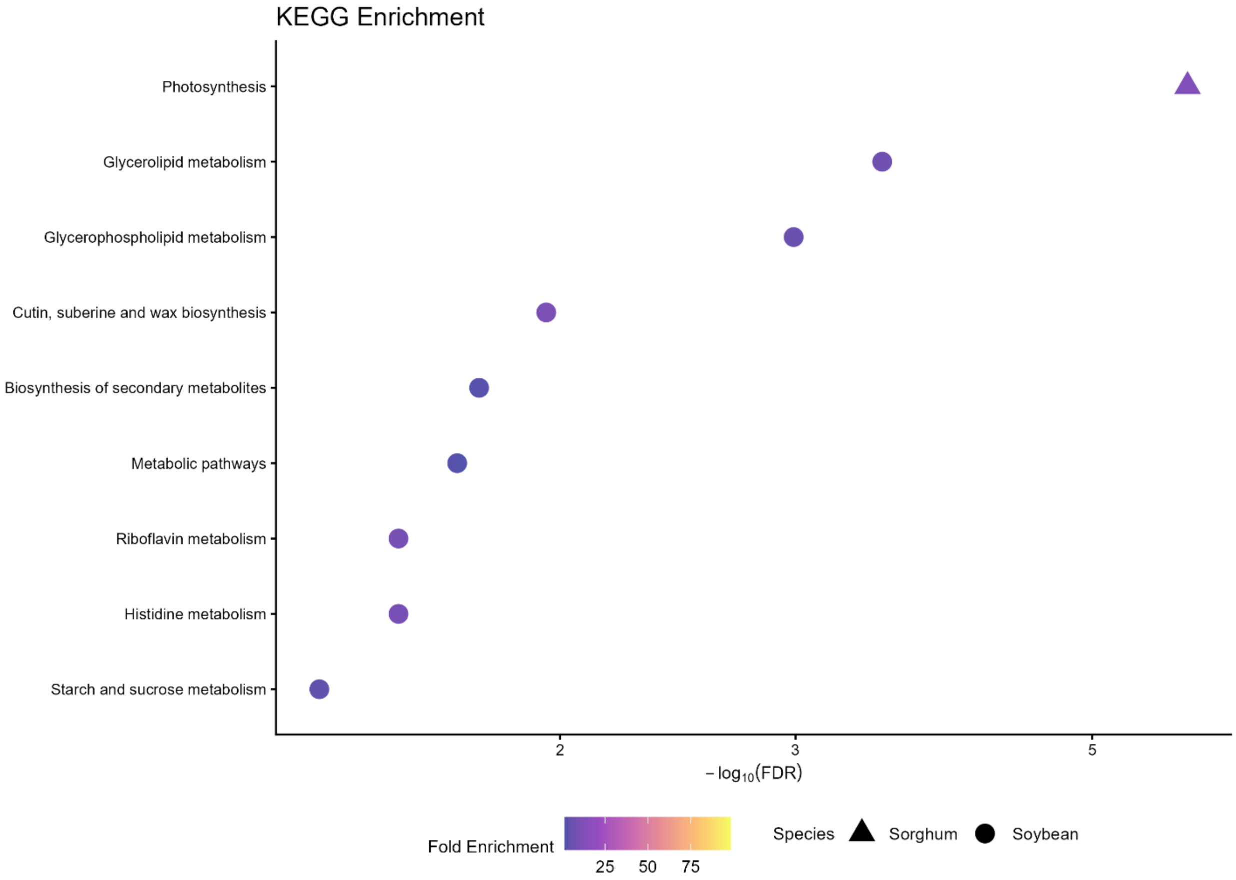


**Supplementary Figure S3.** Top ≤10 significantly enriched (FDR < 0.05) KEGG terms for TWAS-significant genes for each maize, sorghum, and soybean. Filled circle next to term represents significant enrichment across all three species, open circle represents term enriched in at least two species.

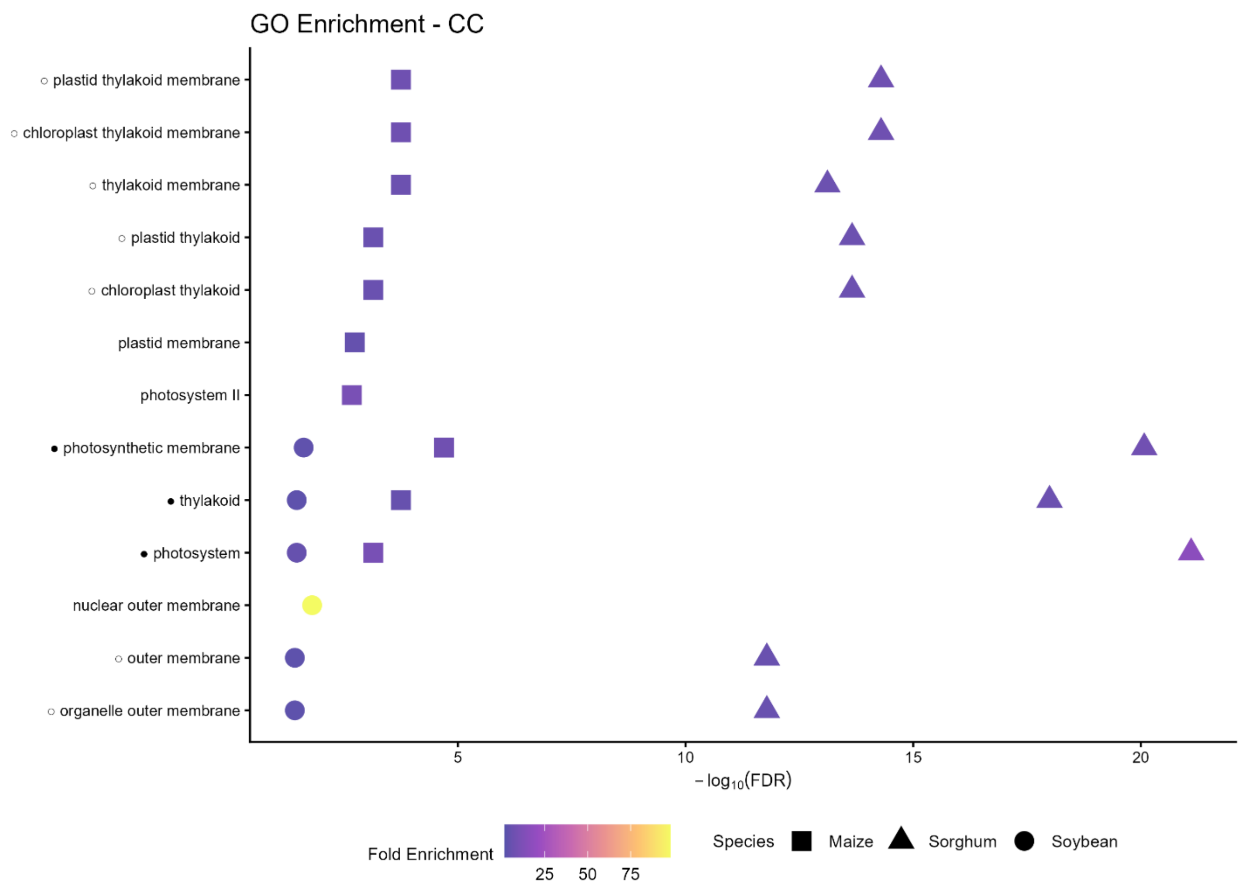


**Supplementary Figure S4.** Top ≤10 significantly enriched (FDR < 0.05) GO-CC terms for TWAS-significant genes for each maize, sorghum, and soybean. Filled circle next to term represents significant enrichment across all three species, open circle represents term enriched in at least two species.

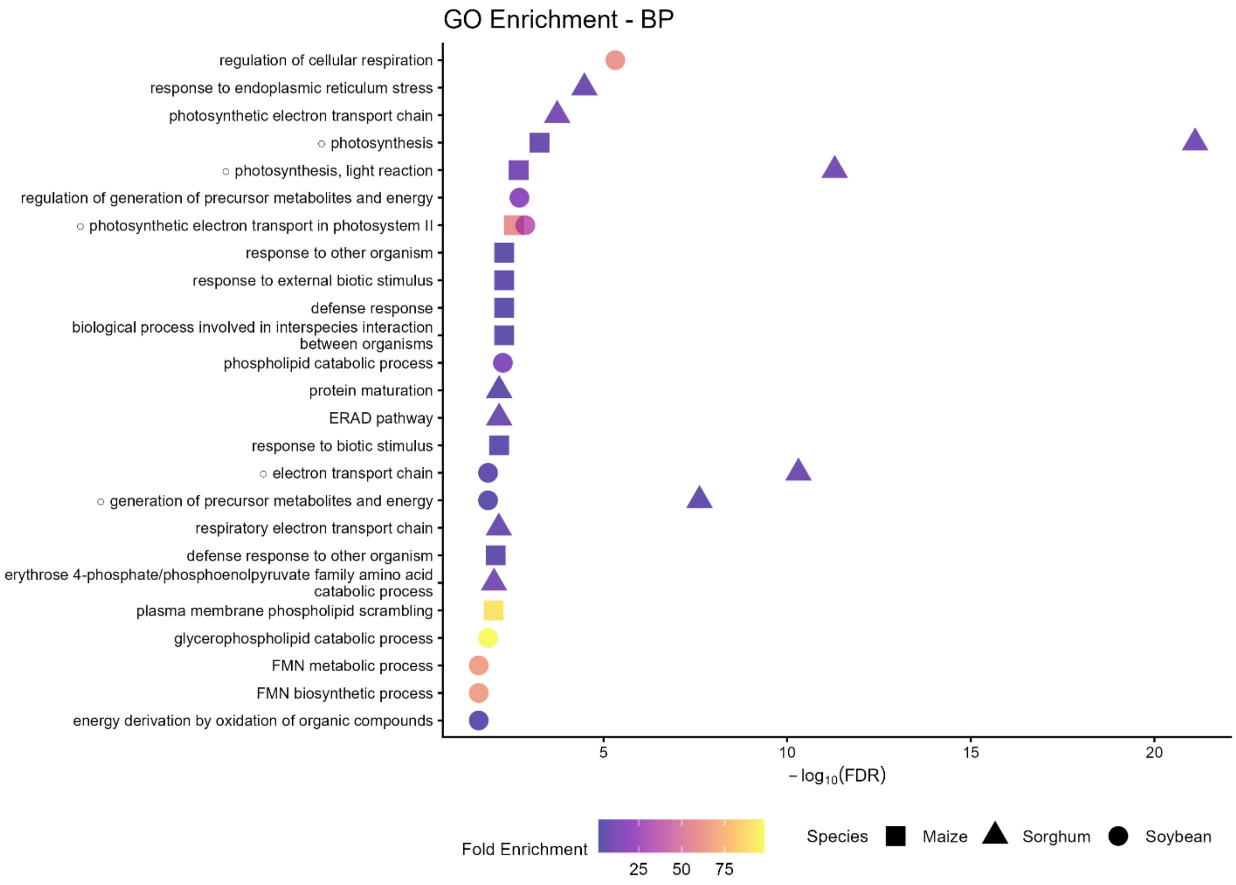


**Supplementary Figure S5.** Top ≤10 significantly enriched (FDR < 0.05) GO-BP terms for TWAS-significant genes for each maize, sorghum, and soybean. Filled circle next to term represents significant enrichment across all three species, open circle represents term enriched in at least two species.

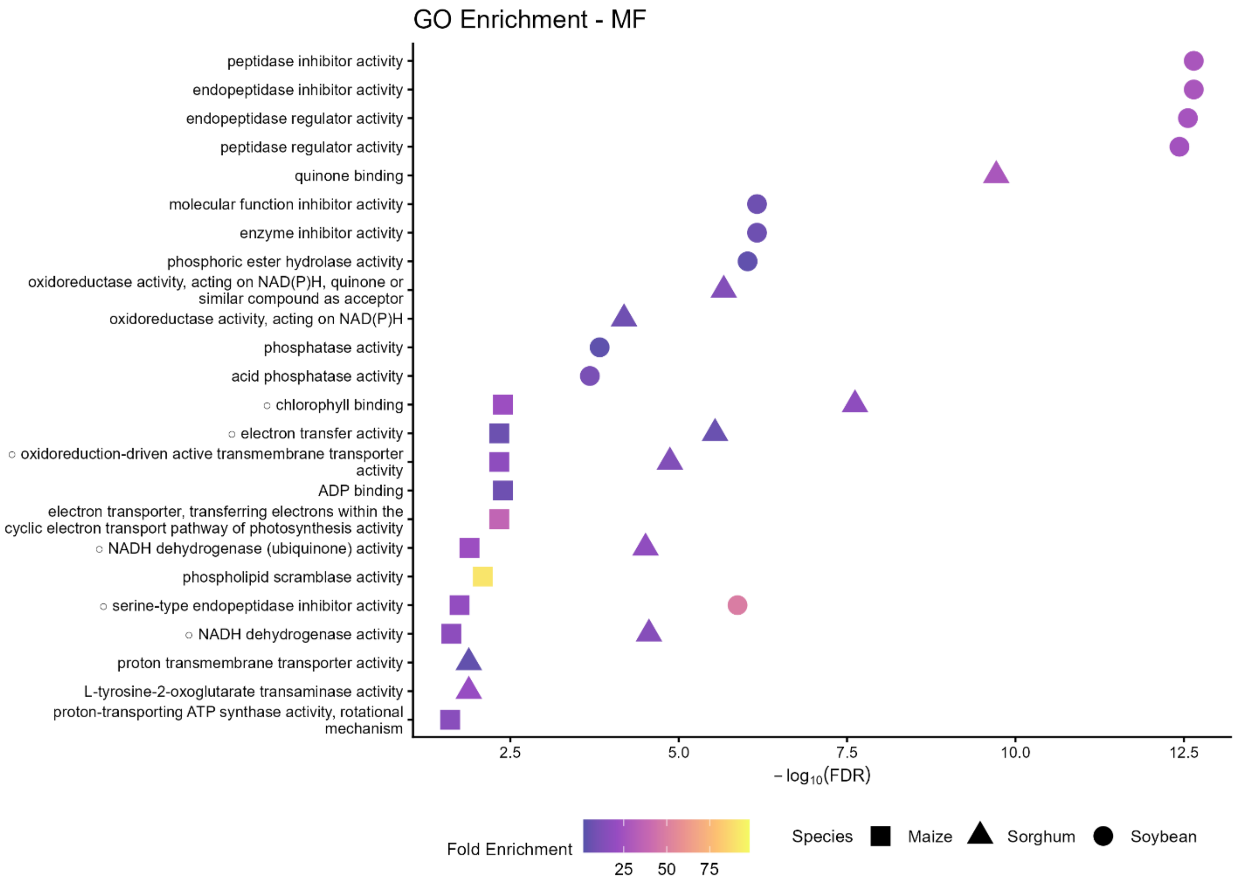


**Supplementary Figure S6.** Top ≤10 significantly enriched (FDR < 0.05) GO-MF terms for TWAS-significant genes for each maize, sorghum, and soybean. Filled circle next to term represents significant enrichment across all three species, open circle represents term enriched in at least two species.

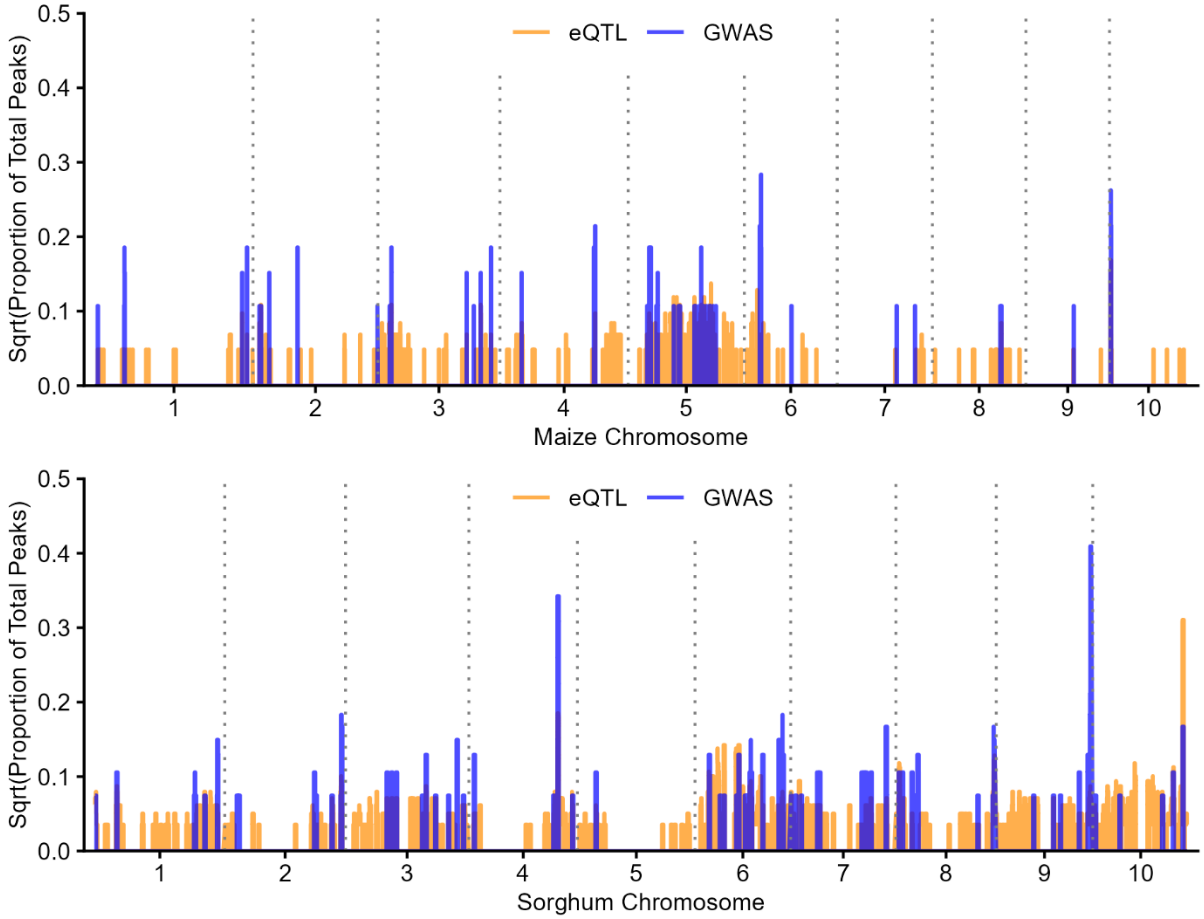


**Supplementary Figure S7.** Genome-wide distribution of fungal abundance–associated regions identified via GWAS or TWAS-eQTLs across maize and sorghum genomes, shown as the proportion of total GWAS or eQTL peaks per species. Proportions were square-root transformed to improve visualization of regions with lower peak densities. peaks were summarized in 1 Mb sliding windows with a 100 bp step.
