## Supplementary material for "Population-scale transcriptomics reveals host genetic control of phyllosphere fungal communities": List of Supplementary Materials

### Supplementary Figures:

**Supplementary Figure S1.** Distribution of fungal read counts per sample across datasets.

**Supplementary Figure S2.** Overlap of fungal taxa detected across maize, sorghum, and soybean for each taxonomic level.

**Supplementary Figure S3.** Top ≤10 significantly enriched (FDR < 0.05) KEGG terms for TWAS-significant genes for each maize, sorghum, and soybean.

**Supplementary Figure S4.** Top ≤10 significantly enriched (FDR < 0.05) GO-CC terms for TWAS-significant genes for each maize, sorghum, and soybean.

**Supplementary Figure S5.** Top ≤10 significantly enriched (FDR < 0.05) GO-BP terms for TWAS-significant genes for each maize, sorghum, and soybean.

**Supplementary Figure S6.** Top ≤10 significantly enriched (FDR < 0.05) GO-MF terms for TWAS-significant genes for each maize, sorghum, and soybean.

**Supplementary Figure S7.** Genome-wide distribution of fungal abundance–associated regions identified via GWAS or TWAS-eQTLs across maize and sorghum genomes, shown as the proportion of total GWAS or eQTL peaks per species.

### Supplementary Tables:

**Provided as single .xlsx file*

**Supplementary Table S1.** Number of fungal taxa retained in each host dataset across taxonomic levels following filtering thresholds.

**Supplementary Table S2.** Number of genetic markers, effective markers, and adjusted significance thresholds for each variant dataset used in association analyses.

**Supplementary Table S3.** Narrow-sense heritability ($h^2$) estimates for fungal relative abundance traits in maize and sorghum.

**Supplementary Table S4.** Significant TWAS associations for fungal relative abundance traits in maize, sorghum, and soybean.

**Supplementary Table S5.** Significant GWAS peaks associated with fungal relative abundance traits in maize and sorghum.

**Supplementary Table S6.** Significant eQTL peaks for TWAS-identified genes associated with fungal relative abundance traits in maize and sorghum.

**Supplementary Table S7.** GWAS and eQTL peaks within the sorghum chromosome 4 hotspot region (Chr04: 57.99–59.96 Mb).

**Supplementary Table S8.** Fungal taxa significantly associated with expression of the sorghum leucine-rich repeat receptor-like kinase gene *Sobic.004G214900*.

**Supplementary Table S9.** GWAS and eQTL peaks within the sorghum chromosome 9 GWAS hotspot region (Chr09: 61.62–63.28 Mb).

**Supplementary Table S10.** GWAS and eQTL peaks within the sorghum chromosome 10 eQTL hotspot region (Chr10: 58.88–60.74 Mb).

### Supplementary Datasets:

**Supplementary Dataset S1.** Maize fungal relative abundance matrix. Includes all samples passing the 2500 fungal-read threshold prior to winsorization and representative-sample selection for genotype-level GWAS/TWAS analyses.

**Supplementary Dataset S2.** Sorghum fungal relative abundance matrix. Includes all samples passing the 2500 fungal-read threshold prior to winsorization and representative-sample selection for genotype-level GWAS/TWAS analyses.

**Supplementary Dataset S3.** Soybean fungal relative abundance matrix. Includes all samples passing the 2500 fungal-read threshold prior to winsorization and representative-sample selection for genotype-level GWAS/TWAS analyses.

**Supplementary Dataset S4.** Orthogroup assignments for TWAS-significant genes identified in maize, sorghum, and soybean.
